# Burrow disturbance, renovation, and collapse in a cavity-nesting seabird

**DOI:** 10.64898/2026.10.05.756071

**Authors:** Luke W. Robinson, Patricia L. Jones, Robert A. Mauck, Jennifer Grady, Liam U. Taylor

**Affiliations:** Department of Biology, Bowdoin College, Brunswick, Maine 04011, USA; Biology Department, Kenyon College, Gambier, Ohio 43022, USA; Cummings School of Veterinary Medicine, Department of Small Animal Clinical Sciences, Tufts University, North Grafton, Massachusetts 01536, USA

**Keywords:** Animal architecture, long-term monitoring, nest site fidelity, researcher disturbance, storm-petrel, seabird conservation

## Abstract

Researcher disturbance can alter the behavior, survival, and breeding success of wildlife, with effects that are especially well-documented in waterbirds. Anecdotal reports suggest that disturbance can even cause cavity-nesting species to alter the structure of their nests. We used a long-term study of a burrowing seabird, the Leach’s Storm-Petrel (*Hydrobates leucorhous*), to ask whether storm-petrels renovate existing burrows and whether such renovation is related to a history of researcher disturbance. Our study was conducted at the Bowdoin Scientific Station on Kent Island, New Brunswick, Canada, where in 2024 we measured the burrow architecture of 363 burrows, including length and structure for occupied and unoccupied burrows both inside and outside a long-term study site with consistent research activity. Of these, 150 burrows had been measured in 2006, allowing for direct longitudinal comparison of burrow architecture. Nearly all burrows changed in length over the 18-year study period (mean absolute change = 20.8 cm ± 18.2 SD). A linear model revealed two contrasting processes of burrow length change: more frequently occupied burrows grew longer, whereas initially longer burrows shortened towards mean burrow length (∼50 cm) through collapse. Occupied burrows inside the long-term study site were on average 34% longer and significantly more complex than burrows adjacent to the site, suggesting that researcher disturbance directly or indirectly reshapes burrows. Overall, these results show that storm-petrel burrows are shaped by cycles of renovation and subsequent collapse that may be exacerbated by human disturbance. Given ongoing population declines in storm-petrels and other cavity-nesting seabirds, this study emphasizes the need to balance scientific research with minimizing disturbance, particularly in long-term studies.

## INTRODUCTION

Understanding the impacts of researcher disturbance is critical for identifying biases in ecological data and minimizing undue stress on animals (Götmark 1992, Soulsbury *et al*. 2020). Disturbance effects are especially well-studied in waterbirds, largely because their tendency to nest on the ground in large, dense colonies makes them readily accessible to investigators.

Population surveys, nest monitoring, and individual tagging can all alter behavior, movement, survival, and breeding success (e.g., Carney and Sydeman 1999, Gauthier-Clerc *et al*. 2004, Carey 2009, Thaxter *et al*. 2016). The consequences of researcher disturbance can vary considerably across waterbird species, populations, or studies, from complete tolerance to injury and reproductive abandonment (Carney and Sydeman 1999, Nisbet 2000).

Among waterbirds, researcher disturbance is particularly relevant to procellariiform seabirds (i.e., albatrosses, shearwaters, petrels, and storm-petrels) for two reasons. First, most procellariiform species are globally imperiled (Dias *et al*. 2019), meaning that special care must be taken to minimize negative effects on their populations. Second, many procellariiforms are nocturnal and burrow-nesting (Warham 1990), making it difficult to study their populations without directly disturbing their nests. Carey (2009) reviews the negative effects of researcher disturbance on procellariiforms, which can include increased nest switching and mate switching, altered hormonal levels, reduced survival, and decreased breeding success.

In his studies of Leach’s Storm-Petrels (*Hydrobates leucorhous*), Gross (1935) suggests another consequence of disturbance: burrow renovation. He anecdotally states that Leach’s Storm-Petrels excavate deeper nest cavities when disturbed during incubation. Burrows and burrow structure play fundamental roles in the survival and reproduction of Leach’s Storm-Petrels by protecting both parents and offspring from predators, heat loss, and inclement weather (Whittow *et al*. 1987, Fricke *et al*. 2015, Richards *et al*. 2024). Indeed, many storm-petrels show high burrow fidelity across years or even decades (Morse and Kress 1984, Blackmer *et al*. 2004, Fricke *et al*. 2015). In turn, excavating new nests or renovating old ones may incur energetic costs both directly, such as through the effort of digging, and indirectly, including by altering the thermodynamic properties of the nest chamber (Whittow *et al*. 1987, Soler et al. 1998, Mainwaring and Hartley 2013, Fricke *et al*. 2015, Richards *et al*. 2024). Aside from general reports about burrows changing length, there is limited information about how storm-petrels build or renovate burrows (Grubb 1970, Pollet *et al*. 2021), let alone the effects of disturbance on those processes.

Here, we used a long-term study of Leach’s Storm-Petrels at Kent Island, New Brunswick, Canada (Fig. 1A) to ask (1) whether storm-petrels renovate existing burrows and (2) whether such renovation is related to a history of researcher disturbance. We measured changes in burrow length and branching structure over an 18-year period and associated these changes with the history of burrow occupancy. We further compared the length and layout of disturbed burrows in the study site with undisturbed burrows adjacent to the site. Our analysis suggests that storm-petrels renovate existing burrows and points towards a nuanced relationship between disturbance, reproductive activity, burrow renovation, and burrow collapse.

**Figure 1.**
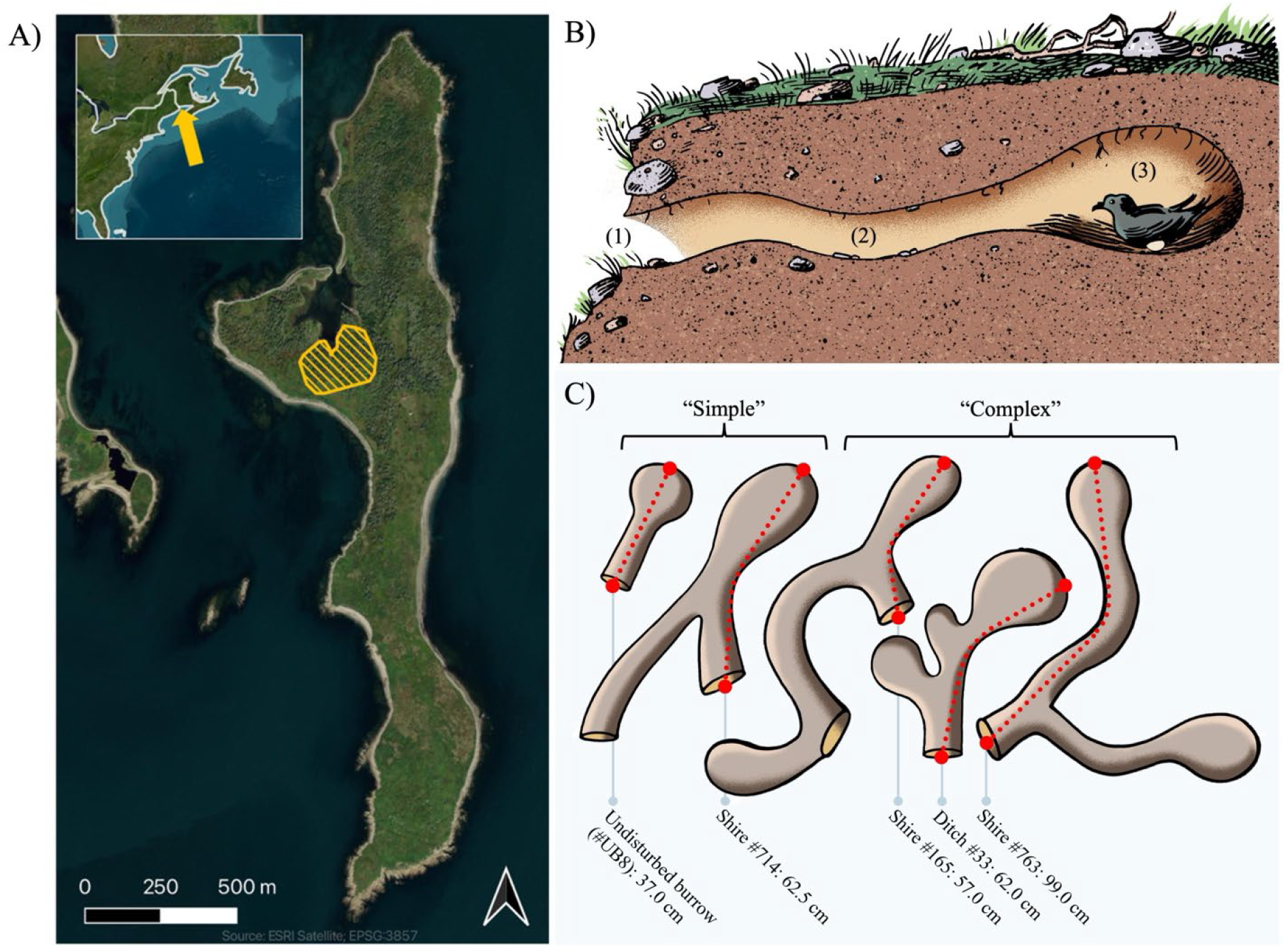
Study site and system. **(A)** The Leach’s Storm-Petrel study area (hatched yellow) on Kent Island, New Brunswick, Canada (44.5819° N, 66.7559° W). **(B)** Cross section of a storm-petrel burrow, including the [1] entrance, [2] tunnel, and [3] nest chamber. **(C)** Example layouts of “simple” and “complex” burrows. Dotted red lines represent the length measurement of each respective burrow. Illustrations by Tim Robinson.

## METHODS

### Study Species

The Leach’s Storm-Petrel (Procellariiformes: Hydrobatidae) is a small, nocturnal seabird with a widely distributed breeding range in the North Atlantic and North Pacific (Pollet *et al*.

2021). Like other storm-petrels, these birds nest in colonies on offshore islands free from terrestrial predators (Harris 1974, Pollet *et al*. 2021). They excavate burrow nests in soft soil, often dug under vegetation or other supportive structures including branches, roots, and stones (Fig. 1B; Fricke *et al*. 2015, Pollet and Shutler 2019). The species exhibits >90% annual mate fidelity and nest site fidelity, with pair bonds documented to last up to 26 years (Morse and Kress 1984, Pollet *et al*. 2026). In the Northwestern Atlantic, Leach’s Storm-Petrel breeding begins at the colony in late April or early May, with egg laying in June and fledging from mid-September to early November. A breeding pair produces a single egg each year, and both parents share the task of incubating the egg and raising the chick over several months (Gross 1935, Pollet *et al*. 2021). Leach’s Storm-Petrels are classified as IUCN Vulnerable, with global population declines potentially linked to increased predation risks, light pollution, and shifting prey distributions at sea (Pollet *et al*. 2023).

### Study Site

Our study was conducted at the Bowdoin Scientific Station on Kent Island, New Brunswick, Canada (Fig. 1A). Located roughly 30 km from mainland New Brunswick, the ∼100 ha island is home to several species of breeding seabirds, including Leach’s Storm-Petrels, American Herring Gulls (*Larus smithsonianus*), Great Black-backed Gulls (*Larus marinus*), Common Eiders (*Somateria mollissima*), and Black Guillemots (*Cepphus grylle*). The Leach’s Storm-Petrel population on Kent Island is estimated at 21,500 breeding pairs as of 2018 (d’Entremont *et al*. 2020). This population has declined by an estimated 1.8% per year since 2000, similar to declines observed across the species range (d’Entremont *et al*. 2020).

Leach’s Storm-Petrels nest throughout Kent Island. The northern half of the island is a combination of dense forest and open sections of ferns, while the southern half is mostly open field habitat. The only other burrowing species on the island is the Muskrat (*Ondatra zibethicus*), whose burrows are easily distinguished from storm-petrel burrows by size and moisture. In recent years, a small number of predatory North American River Otters (*Lontra canadensis*) have also been observed on Kent Island.

Data were collected as part of a long-term study on Kent Island at two sub-sites (the “Shire” and the “Ditch”; Fig. 1A). Burrows in the Shire have been monitored every year since 2003, while burrows in the Ditch have been monitored, with gaps in research, since 2008.

Burrows in these sub-sites are marked with unique ID tags and monitored during the breeding season. Monitoring begins in June, before most eggs are laid, and continues through July when most chicks have hatched.

Every burrow is checked multiple times each year using established protocols to determine egg lay date, hatch date, and identity of breeding parents (Mauck *et al*. 2004). These protocols involve “grubbing” — a researcher inserts their arm down a burrow tunnel, feels the contents of the nest chamber, and, if necessary to collect data, removes the egg and removes and identifies any parents that are present. If the nest chamber cannot be reached, access holes are dug along the burrow tunnel, taking care to preserve the integrity of the burrow, and then resealed with a weighted wooden cover. Adult birds are banded with a metal leg band or identified by existing bands, and morphometrics (weight, tarsus, and wing chord) and egg measurements (length, width) are recorded. Birds and eggs are then returned to the nest chamber. Thus, within the study site, each occupied nest is entered at least four times per season: once for egg discovery, two more times for identifying each parent, and a fourth time to check for hatching. Birds may experience additional disturbance, including blood samples for genetic analyses, or additional nest visits to collect complete data, as it is difficult to predict the timing of egg laying, partner changes, and egg development.

### Burrow Measurements

In 2024, we measured the architecture (i.e., length and/or branching structure) of occupied burrows (*n* = 136) and unoccupied burrows (*n* = 185) within the study site. Occupancy was defined by the presence of an egg at any time during the breeding season. Some of these burrows (*n* = 150) were previously measured in 2006 in the Shire, allowing us to assess changes in architecture following 18 additional years of researcher disturbance. Burrows inside the study site were opportunistically selected for measurement during regular nest checks, additionally targeting burrows that had been previously measured in 2006. This sample was supplemented with a random set of unoccupied burrows within the study site. We did not measure intertwined burrows with multiple active nest sites (i.e., more than one breeding pair sharing a set of burrow entrances). As a result, our sample excludes some of the longest and most complex burrows in the study site.

In 2024, we also randomly selected 50 potential burrows outside the study site for measurement, retaining only those that could be confirmed as burrows because they were active with an egg (*n* = 42). These burrows were located directly adjacent to the study site and to our knowledge have never been disturbed. The habitat in this undisturbed area is identical to within the study site, differing only in its disturbance history. Data from outside of the study site was limited to occupied burrows because, in the absence of previous identification, we could not sample a representative distribution of unoccupied or collapsed burrows.

Burrow length was measured as the total length from the burrow entrance to the farthest extent of the burrow, behind the nest chamber (Fig. 1B–C). If a burrow had multiple entrances or nest chambers, we measured from the entrance nearest to the longest nest chamber (Fig. 1C).

Measurements were taken with twine, staked at the furthest extent from the entrance, and pulled taut. The twine was pinched directly underneath the most receded part of the roof of the burrow entrance, then the stake was removed, and the twine was measured. All 2024 burrow lengths were rounded to the nearest centimeter to match the precision of 2006 measurements. For burrow structure, we counted the number of entrances and tunnels within a burrow. Entrances were only counted if they provided access to the nest chamber. We categorized burrow structure into two categories: simple (no branching), or complex (branching to multiple potential nest chambers, although all of these burrows contained at most one egg; Fig. 1C). Long-term monitoring provided data about the occupancy of burrows from 2006 to 2024.

### Data Analysis

All analyses were conducted in R version 4.6.0 with the *tidyverse* packages (Wickham *et al*. 2019, R Core Team 2026). All tests were two-tailed. Significance was assessed at *α* = 0.05. Due to multiple measurements and comparisons, different statistical analyses used different subsets of measured burrows; we provide sample sizes for each test separately.

We assessed burrow renovation by comparing burrow lengths between 2006 and 2024 using a paired t-test (*n* = 150). We assessed burrow structural changes between 2006 and 2024 using McNemar’s test on pair proportions, using the same set of burrows measured in both years (*n* = 150).

To investigate the mechanisms of length change in these burrows, we then fit a linear model with two fixed effects: (1) the number of years that breeding pairs actively occupied each burrow from 2006 to 2024 and (2) the initial length of the burrow in 2006. These variables were chosen to test whether (1) more frequently occupied burrows experienced more renovation, and (2) overall burrow size influenced the likelihood of subsequent maintenance or structural collapse. This analysis only included burrows measured in both 2006 and 2024 (*n* = 150), which were all from the Shire sub-site with consistent information about annual occupancy.

We compared the lengths of occupied disturbed (i.e., within the study site) and undisturbed (i.e., adjacent to the study site) burrows using independent-sample t-tests (occupied disturbed: *n* = 134; occupied undisturbed: *n* = 42). Differences in the distribution of simple and complex burrows between disturbed and undisturbed areas were assessed with a Pearson’s chi-squared test (occupied disturbed: *n* = 136; occupied undisturbed: *n* = 42).

## RESULTS

Burrow architecture was highly dynamic over the 18-year study period, with nearly every burrow either increasing or decreasing in length (absolute change in burrow length, mean ± SD: 20.8 ± 18.2 cm; Fig. 2). Occupancy of burrows included in the longitudinal study ranged from 0 years to 18 years, with a mean occupancy of 7.1 ± 5.3 years. Changes in burrow length occurred in both directions and were often substantial. Increases in burrow length over 18 years (*n* = 69 burrows) ranged from 1 to 57 cm, while decreases (*n* = 79 burrows) ranged from -1 to -82 cm. Looking across all burrows, mean length decreased from 57.9 cm (± 15.8) in 2006 to 52.3 (± 24.3) in 2024, a statistically significant decline (paired t-test, *t* = 2.52, *df* = 149, *p* = 0.01). This overall decline includes 8 burrows measured in 2006 that completely collapsed by 2024 (i.e., burrow length of 0 cm). The overall directional change in burrow length is not significant if these completely collapsed burrows are excluded (paired t-test, *t =* 1.29, *df* = 141, *p* = 0.20).

**Figure 2.**
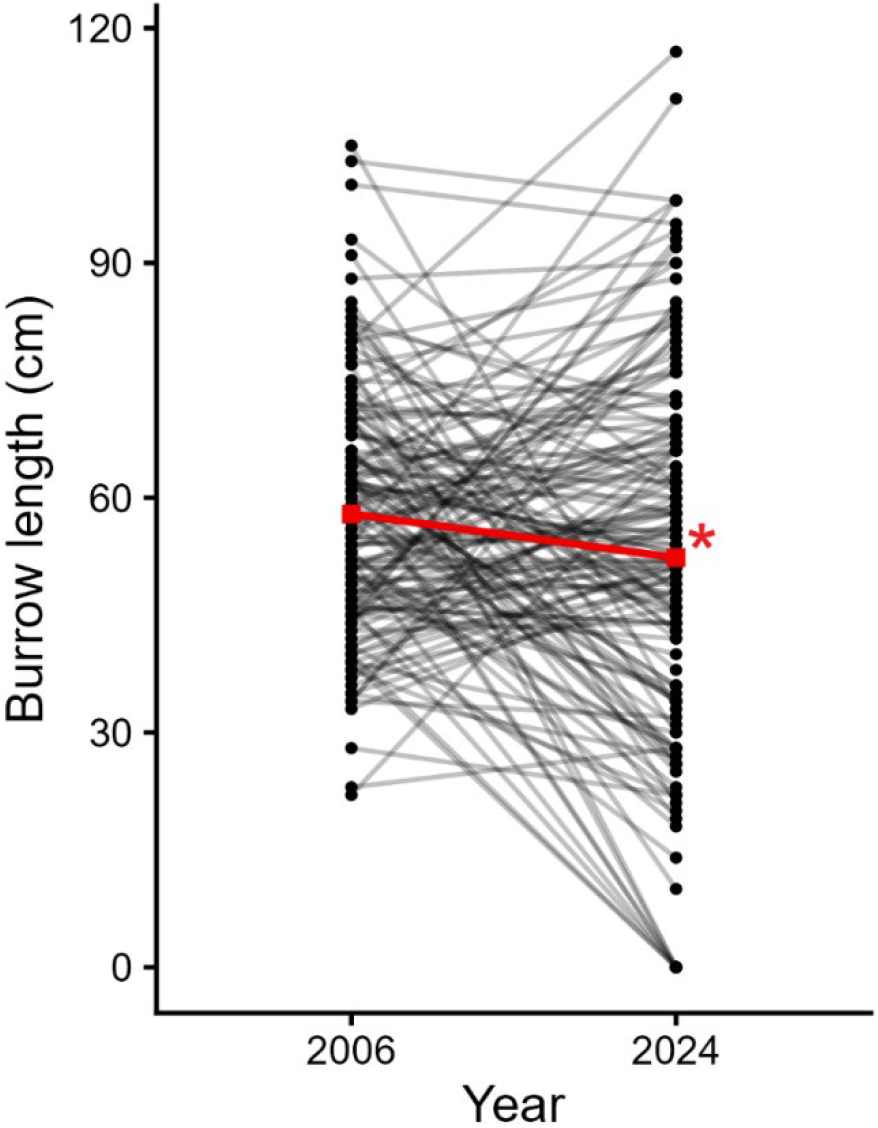
The nesting burrows of Leach’s Storm-Petrel are dynamic, with nearly every burrow changing in length across 18 years of monitoring. Black points indicate individual burrow lengths, with grey lines connecting the same burrow across time points. Red points show annual verages, with a small average decline across the time points (* indicates *p* = 0.01).

Burrows significantly changed in structure across the study period (McNemar’s test, *χ*^2^ = 4.0, *p* = 0.05). Comparing the structures of these burrows across years, 12.7% of measured burrows had complex structure in 2024, versus only 5.3% of burrows in 2006 (Table 1). Eighteen simple burrows in 2006 (12.0%) were categorized as complex in 2024, indicating burrow renovation or merging, while 7 complex burrows in 2006 (4.7%) had collapsed to simple burrows by 2024. The significance of this result is highly sensitive to the small number of complex burrows; removing just one complex burrow from 2024 results in a non-significant test (McNemar’s test, χ^2^ = 3.4, *p* = 0.07).

**Table 1.** Burrow structure within the study site, with the same set of burrows measured in both 2006 and 2024. “Simple” burrows had only one nest site, whereas “complex” burrows branched to multiple nest sites.

|  | 2024: Simple | 2024: Complex |
| --- | --- | --- |
| 2006: Simple | 124 | 18 |
| 2006: Complex | 7 | 1 |

A linear model revealed two contrasting effects on burrow length change: more frequently occupied burrows experienced significantly more growth, whereas initially longer burrows experienced significantly more collapse (*R*^2^ = 0.43, *F*_2,147_ = 56.8, *p* < 0.001; Fig. 3). Burrow growth was positively associated with occupancy, with final length increasing by 2.46 cm per year of recorded occupancy (*β* = 2.46, *t* = 7.65, *p* < 0.001; Fig. 3A), and burrow growth was negatively associated with initial length in 2006, decreasing by 0.88 cm for each additional centimeter of initial length (*t* = -8.26, *p* < 0.001; Fig. 3B).

**Figure 3.**
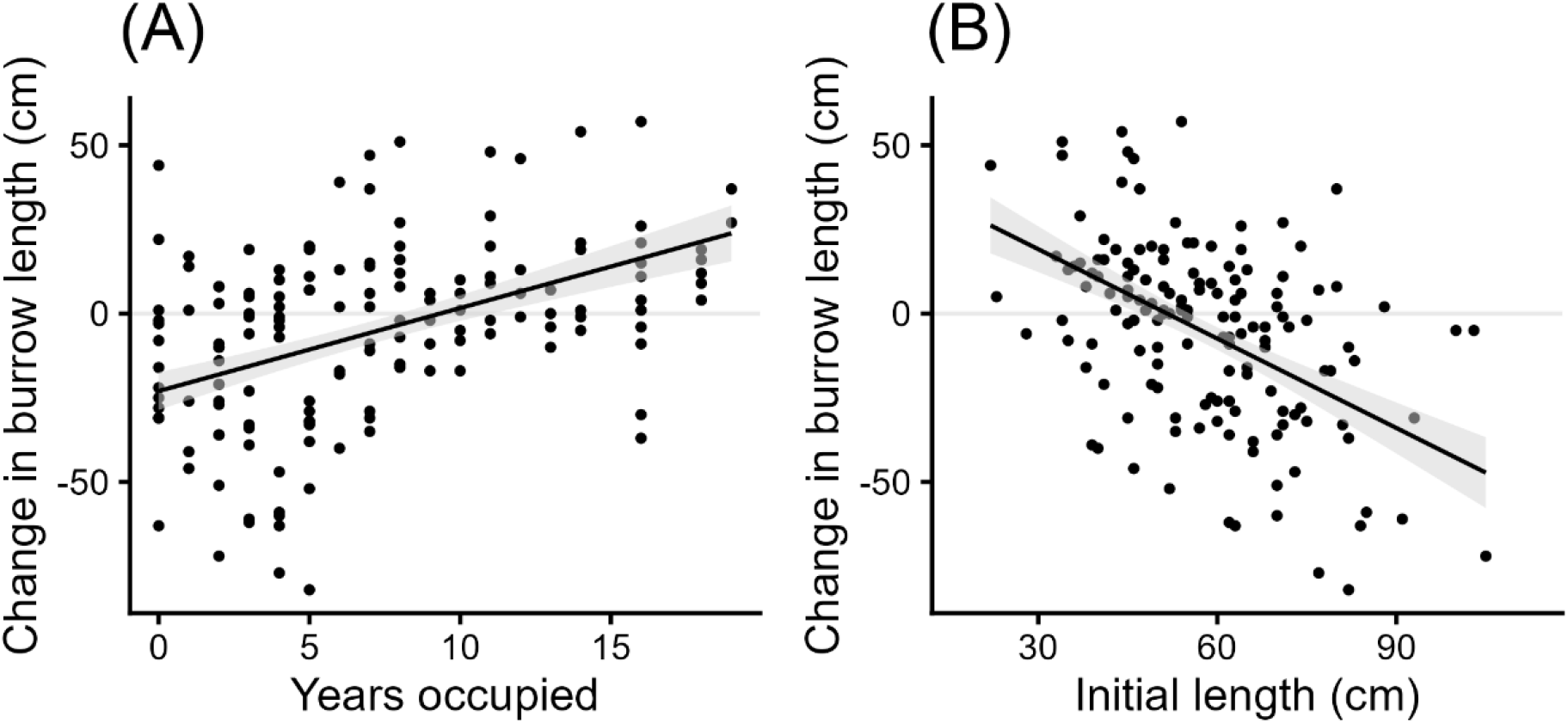
The length of Leach’s Storm-Petrel burrows **(A)** increases with repeated occupancy and **(B)** decreases through partial collapse when they have longer initial lengths. Points represent individual burrows, with change in length calculated across 18 years of monitoring (2006 to 2024). Black lines show partial effects from a multiple linear regression model, with the non-focal predictor held at its mean; shaded areas indicate 95% confidence intervals for the fitted values.

Comparing actively occupied burrows measured in 2024, disturbed burrows inside the study site were significantly longer than undisturbed burrows adjacent to the study site. Among these occupied burrows, mean disturbed burrow length was 70.0 cm (± 19.8 SD, range 36–173 cm), while mean undisturbed burrow length was 52.2 cm (± 14.4 SD, range 28–88; unpaired t-test, *t* = -6.38, *df* = 93.8, *p* < 0.001; Fig. 4). Excluding one especially long disturbed burrow (length = 173 cm) does not change this result (unpaired t-test, *t =* -6.34, *df* = 83.6, *p <* 0.001). Unoccupied burrows within the study site that were not fully collapsed (*n* = 119, mean length = 50.6 cm ± 19.3 SD) were significantly shorter than occupied burrows in the study site (unpaired t-test, *t* = -7.91, *df* = 248.9, *p* < 0.001) but did not significantly differ in length from occupied burrows adjacent to the study site (unpaired t-test, *t =* 0.55, *df =* 95.9, p = 0.59).

**Figure 4.**
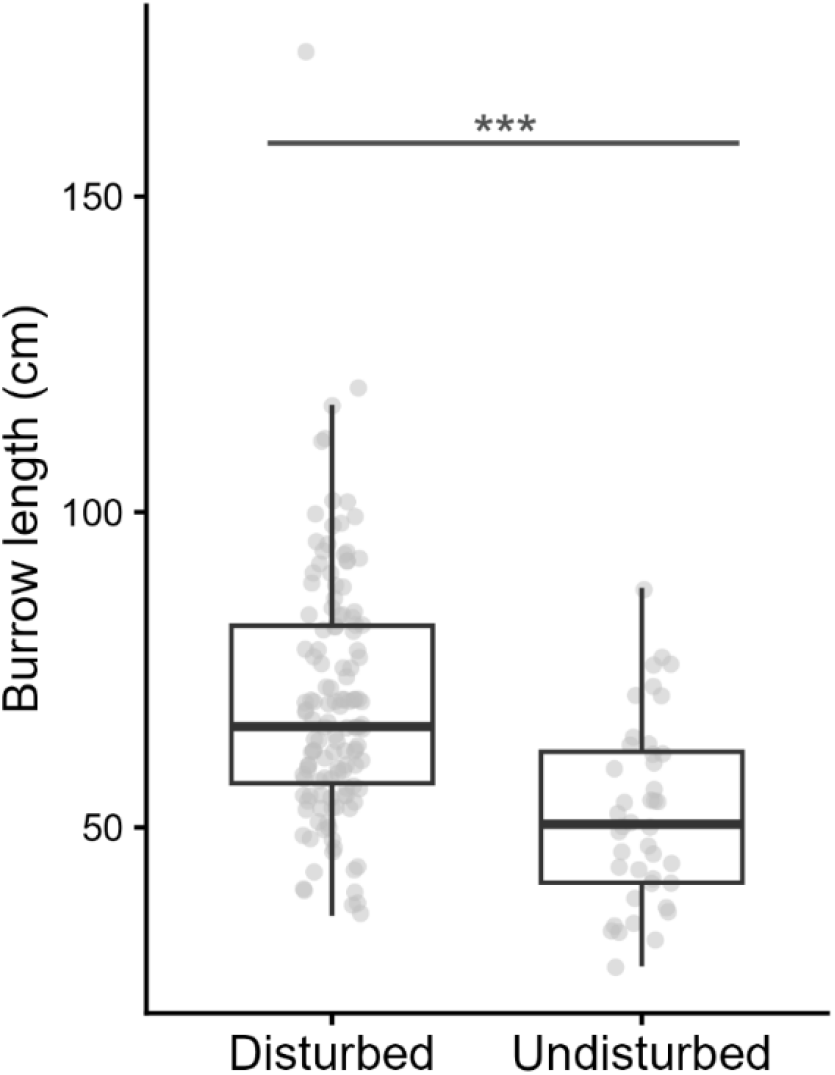
Burrow length in undisturbed areas (i.e., directly adjacent to the study site) and disturbed areas (i.e., inside the study site). Boxes indicate 25^th^–75^th^ percentile, with lines at median and whiskers extending 1.5 times the inter-quartile range (*** indicates *p* < 0.001).

Comparing the structure of disturbed versus undisturbed burrows, 16.2% of burrows (*n =* 22) inside the study site had complex structure compared to only 2.4% of burrows (*n* = 1) outside the study site (Table 2). This difference was statistically significant (Pearson’s chi-squared, χ^2^ = 4.27, *p* = 0.04). This comparison was again limited by the relatively small sample of complex burrows and because exceedingly complex burrows with multiple active nest sites—all of which were located within the disturbed study site—were not included in our 2024 sampling.

**Table 2.**
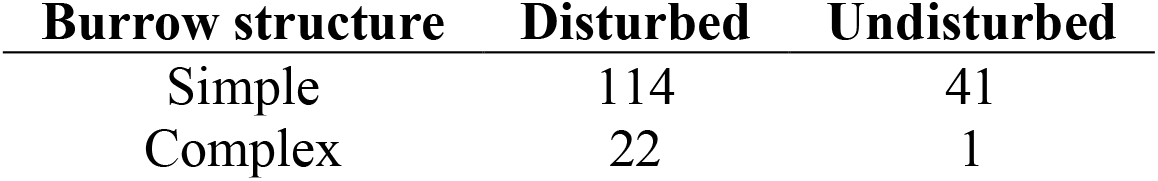
Burrow structure in disturbed areas (i.e., inside the study site) and undisturbed areas (i.e., immediately adjacent to the study site), measured in 2024. This comparison only includes actively occupied burrows.

| Burrow structure | Disturbed | Undisturbed |
| --- | --- | --- |
| Simple | 114 | 41 |
| Complex | 22 | 1 |

## DISCUSSION

We measured changes in the burrows of an imperiled seabird, the Leach’s Storm-Petrel, following an 18-year period of intensive research monitoring. Our first key finding is that nearly all burrows changed length over the study period, including both increases and decreases in length (Fig. 2). Our data confirm that individual nest sites in this species can change dramatically over decades, with the most extreme observed increase in length being 57 cm, and the most extreme decrease in length being 82 cm. We predict that dynamic burrows are common across species that nest in soft substrates (e.g., other storm-petrels, bank-nesting swallows, bank-nesting coraciiforms). Of course, species that nest in hard substrates—such as guillemots or puffins nesting in boulders or cliffs—should have more stable nest cavities.

Our second key finding is that changes in burrow length are likely to arise from two distinct processes: renovation and collapse. Burrows that were more frequently occupied across years tended to increase in length (Fig. 3A). In contrast, initially longer burrows experienced significantly greater rates of collapse (Fig. 3B), some completely collapsing over the course of the study. Assuming that birds do not backfill their burrows, burrows can only decline in length through a process of partial collapse. Longer burrows may collapse more frequently due to structural weaknesses, as more soil weighs on the unsupported portions of longer or deeper tunnels. In addition, breeding birds’ preferences for burrows with specific depths, angles, or temperatures (Fricke *et al*. 2015) may mean that individuals do not actively maintain the full length of the longest burrows, resulting in regression towards a mean preferred burrow size.

More frequently occupied burrows could increase in length because, in hosting the birds and eggs that researchers actually monitor, they experienced more frequent and direct disturbance. Indeed, our third key finding is that disturbed burrows inside the study site were significantly longer than undisturbed burrows adjacent to the study site. Looking across occupied burrows, the mean length of disturbed burrows was 18 cm longer (+34%) than undisturbed burrows (Fig. 4). Two possible mechanisms for this observation are (1) disturbance compels existing occupants to dig deeper to avoid perceived threats (Gross 1935), or (2) disturbance increases parent turnover (Blackmer *et al*. 2004) with new occupants excavating upon arrival.

In addition to burrow length, disturbance may also affect burrow structure either directly (e.g., from temporary access holes created by researchers) or indirectly (e.g., disturbance-induced renovation by the birds). Indeed, while complex burrows within the study site are commonly merged from multiple entrances within the soil, the only complex burrow outside the study site was in the base of an upturned tree, creating natural intertwined cavities in the roots. The most labyrinthine burrows inside the study site were excluded from the present study because their length could not be reliably determined, and because they often contained multiple active nest sites attended by different mating pairs. The combined tunnel length of one of these burrows likely stretches many meters.

Although Leach’s Storm-Petrels prefer longer burrows (Fricke *et al*. 2015), longer burrows may also be less structurally stable. Complete burrow collapse would necessitate switching burrows, which is often coupled with partner change in this species (Morse and Kress 1984, Blackmer *et al*. 2004, Mariné and Bernard 2020). Both mate and nest fidelity may help procellariiforms better coordinate incubation schedules and foraging routines across multiple years of reproduction (Bried *et al*. 2003). Consequently, new breeding pairs are more likely to fail than pairs that have bred together for multiple seasons (Weimerskirch 1990). Thus, we expect burrow collapse to have real reproductive consequences in this species.

More generally, our results suggest a tradeoff in nest site architecture: longer burrows may confer thermoregulatory benefits by decreasing convective heat loss during incubation and brooding (Fricke *et al*. 2015), but they require energetic investment to excavate (Grubb 1970, Mainwaring and Hartley 2013) and are more prone to collapse or less likely to be maintained (Fig. 3B). Based on predictions from the multiple linear regression with years of occupancy held at its average (Fig. 3B), ∼52 cm may represent a stable optimum for burrows in this study site.

Longer burrows are predicted to collapse towards this size, whereas shorter burrows are predicted to lengthen. This value matches the mean length of undisturbed burrows reported in this study (52.2 cm ± 14.4 SD, *n* = 42), aligns with the average length of burrows on Kent Island reported by Gross in 1935 (50.8 cm, range 28–87 cm, *n* = 40), and falls within the range of average burrow lengths reported from minimally disturbed sites outside Kent Island (43–74.9 cm; Pollet *et al*. 2021, Vermeer *et al*. 1988). We hypothesize that differences in the thermodynamics and structural stability of substrates can shift the stable burrow size across breeding colonies.

Overall, our results suggest that storm-petrel burrows are structurally dynamic and reshaped with disturbance. This study is limited in that it represents only two ‘snapshots’ in time. Continuous monitoring of burrows throughout their lifespan will reveal additional insights into the dynamics of burrow architecture changes. The energetic and reproductive consequences of burrow renovation and collapse might compound other disturbance effects, such as increased mate switching (Blackmer *et al*. 2004, Carey 2009), to ultimately impact reproductive success.

As an IUCN Vulnerable species, Leach’s Storm-Petrels have experienced substantial population declines attributed to elevated predation rates and artificial light attraction (Pollet *et al*. 2023). Seabirds in general are threatened both locally and globally by introduced predators, fisheries, disease, and climate change (Dias *et al*. 2019). While we predict that the disturbance effects on burrow architecture are relatively minor, more akin to disturbance effects on behavior (Nisbet 2000) than direct mortality (Gauthier-Clerc *et al*. 2004), our findings serve as a reminder of the unexpected consequences that disturbance can have on seabirds’ lives.

Long-term studies are critical for seabirds because their population dynamics play out over decades (Nisbet 1989, 2025). We thus renew the call for long-term studies that take care to minimize disturbance, protecting both avian populations and the insights of ecological data.

Modern technologies, such as automatic scales and radio-frequency identification (RFID) tags, can provide high-resolution data, record undisturbed behaviors, and limit disturbance (e.g., Hentati-Sundberg et al. 2025, Mauck et al. 2025). Continuing to integrate these technologies with classic monitoring approaches promises to increase the value of long-term studies while reducing their impact on sensitive species.

## ACKNOWLEDGEMENTS

We thank the students, faculty, and staff on Kent Island for their work in the long-term study of storm-petrels. We are especially grateful to Ian Kyle and Riley Simon for their assistance during fieldwork. Long-term monitoring of Leach’s Storm-Petrels on Kent Island was conducted under Canadian Wildlife Service Permit #02109.

## AI DECLARATION

We used ChatGPT (GPT-4) and Claude (Sonnet 4) to generate portions of regression and visualization code and to recommend sentence-level manuscript edits for clarity. We used Claude (Opus 5.5) to conduct a final verification of the codebase and flag typographical and syntax errors in the manuscript.

